# Identifying cooperative genes causing cancer progression with dynamic causal inference

**DOI:** 10.1101/2023.11.22.568367

**Authors:** Andres M. Cifuentes-Bernal, Lin Liu, Jiuyong Li, Thuc Duy Le

**Author notes:** Corresponding authors: andres.cifuentes.

## Abstract

Cancer progression is driven by complex gene interactions, which are not fully understood due to the dynamic and multifaceted nature of the disease. Traditional methods for identifying genes driving cancer typically focus on detecting mutational events, often neglecting other genetic alterations and the temporal dynamics of the cancer development. This study introduces a novel approach that expands our understanding of cancer by identifying cooperative networks of cancer drivers through dynamic causal inference, complementing classical mutation-based methods.

We developed a data-driven causal inference method that integrates a temporal dimension to gene expression data through pseudotime analysis. By modelling cancer progression as a dynamic system, the method evaluates gene interactions and their causal relationships using causal kinetic models (CKMs). This approach enables detecting cancer driver genes regardless of their mutational status. The approach was applied to both single cell and bulk sequencing datasets of breast cancer to identify and rank cancer driver genes based on their stability on the causal model.

The proposed method effectively identifies and ranks cancer driver genes by considering their dynamic interactions during cancer progression. It systematically identified cancer driver genes in both categories: those with single nucleotide variants (SNVs) and those with other alterations. The method demonstrated a significant overlap with known cancer driver genes from the Cancer Gene Census, validating its effectiveness. Additionally, the method uncovered novel driver genes that act cooperatively to influence cancer development, offering a more comprehensive view of the genetic mechanisms underlying cancer. Our approach, along with the scripts for the experiments and datasets used, can be found at https://github.com/AndresMCB/DynamicCancerDriverKM

## 1 Introduction

Cancer is one of the leading causes of death worldwide, accounting for one in six deaths and nearly 18 million new cases in 2020 [15]. One key aspect of this disease is that treating it in its early stages sharply increases the likelihood of survival [47]. For this reason, understanding the dynamics of cancer (i.e., how it progresses) is crucial for successfully detecting and treating cancer. However, our current knowledge of the dynamics of cancer progression is far from complete, in part due to the challenges involved with the modelling of cancer at different stages [15].

Cancer genesis and development are linked to aberrations in genes commonly known as cancer drivers. Such aberrations have been demonstrated to provide a strategic evolutionary advantage to certain cells by altering critical biological processes in these cells. However, both genetic aberrations and their impacts on biological processes have not been fully characterised. Therefore, detecting cancer drivers and understanding their temporal behaviour are crucial tasks in cancer research. Most of the current approaches for inferring cancer drivers focus on gene mutations within cancer cells that consistently appear at a higher rate than expected by chance [3, 27, 37]. In this paradigm, the discovery of cancer drivers relies on the frequency of the occurrence of such mutations (commonly called driver mutations).

Despite the success of current mutational approaches for discovering cancer drivers, it is now known that aberrations that cause cancer are not limited to mutations and they can appear at different levels, such as the epigenetic level [5, 24, 35]. Moreover, rare alterations (i.e. low-frequency events) contributing to cancer development are difficult to detect in data. As a result, significant effort has been committed to integrating additional information in order to compensate for the incompleteness in the approaches based on mutations. These new methods can be categorised as network-based methods (e.g. [6, 34, 50]) and methods integrating multi-omics (e.g. [26, 38]). Interesting reviews on these methods can be found in [2, 28, 35].

Most current methods for discovering cancer drivers do not consider the temporal information of the disease, nor do they include suitable procedures for testing the causal nature of cancer progression. Consequently, current methods provide few insights into the accurate modelling of cancer as a dynamic system. Furthermore, the dynamics of the causal relationships that occur during cancer progression are neglected. A lack of formal dynamic causal inference procedures results in a reduced capability of discovering cancer drivers from data.

The information required for the dynamic modelling of cancer can be obtained from time series data (e.g. [25, 45]). Time series are formed by collecting data from samples at different points in time during the progression of the disease. In practice, the collection of time series that describe biological processes involved in cancer is difficult and costly. Therefore, most of the available data correspond to snapshots taken from samples of cancer tissues (bulk data). Consequently, the vast majority of cancer driver inference methods have been developed for bulk data, which captures a minimal amount of information on the dynamics of cancer drivers.

One of the most promising approaches for extracting the underlying dynamics of biological processes relies on the *pseudotime* concept [4]. *Pseudotime* inference methods commonly use the gene expression of samples at different stages of a biological transition to calculate a *pseudotime* score. This score aims to reflect the position of the sample in the progression of the biological transition. A large number of studies have shown that *pseudotime* scoring is intrinsically temporal. Furthermore, *pseudotime* has been shown to be useful for extracting part of the underlying dynamics of biological processes, even from bulk data [4, 11, 12, 40, 49].

By definition, a biological process involves the joint action of a specific set of genes, the expressions of which are highly regulated to follow a particular temporal sequence. Individual identification of cancer drivers (which is popular in most current methods) sidesteps this critical fact. To overcome this limitation, in this paper an integrative method that models the causal relationships of cancer drivers influencing cancer progression (groups of driver genes rather than single drivers) is proposed. The method includes a temporal dimension to the data and formal procedures for testing causality in time-dependant data.

In the proposed method, to explore the dynamic aspect of cancer drivers, a temporal dimension to the data is included using the concept of *pseudotime*. Additionally, cancer drivers are identified by assessing the causal influence that driver genes have on the gene expression alterations occurring during cancer development. The proposed method explores the dynamics of cancer and the causal nature of driver genes by modelling the disease as a set of *causal kinetic models* (*CKMs*) [33]. A *CKM* is an extension of the *structural causal models* to the setting of *ordinary differential equations* (*ODEs*). Specifically, *CKMs* are used to model causal relationships involving cancer drivers as a collection of causal *ODEs*. The implementation of the method is discussed in section 2.

The method is applied to the TCGA-BRCA cancer dataset [46] and the single cell RNA sequencing dataset in the NCBI GEO database (accession GSE75688) [10]. The results suggest that the proposed method is capable of systematically identifying cancer driver genes (hyper-geometric test p.value *<* 0.05) based on the dynamics of the interactions detected in the data. Additionally, the proposed method returns the set of dynamic models that are the most consistent with the observed data, providing insights into the underlying behaviour of biological processes in cancer as a dynamic system (in the sense that the system’s current state depends on the past). Furthermore, the proposed method can detect interactions and collaborations among the genes that cause cancer rather than focusing exclusively on the influence of single cancer driver genes on cancer development. As a result, the proposed method can elucidate the behaviour of genes that act together to cause cancer progression.

## 2 Materials and Methods

### 2.1 Problem definition

The premise of the proposed method is that genes driving cancer progression disrupt the balance of core biological processes, such as cellular division and apoptosis, [14, 20, 21]. Because alterations in biological processes due to cancer drivers are progressive over time, the causal relationships involving driver genes can be considered a dynamic system. It is possible to consider each biological process as a dynamic system of genetic interactions. However, cancer itself is a highly complex dynamic system because it progresses over time (i.e. its future states depend on the past).

Such a dynamic system involves both the interaction and the collaboration of genes in the regulation of biological processes. Despite significant advances in genomics, there are still gaps in our understanding of how genes interact with each other. In particular, models of cancer as a dynamic system involving interactions and collaborations between genes have not yet been fully explored. In this paper, a causal model of cancer (as a dynamical system) that takes into account the collaborations and interactions of genes causing cancer progression is presented.

#### 2.1.1 Assumptions

In the proposed method, genes are considered random variables and the data correspond to the gene expression of *m* genes and *n* samples represented by a matrix **G** ∈ ℝ^*n*×*m*^. It is assumed that a subset of genes in **G** causes disruptions in one or more core biological processes during cancer progression. To assess causality, a target variable *y* ∈ ℝ^*n*×1^ is also required. It is expected that the changes in the target variable are due to the influence of cancer drivers. Consequently, the target variable encodes significant information on cancer progression. Suitable target variables for the proposed method include relevant genes for cancer, such as well-recognised biomarker genes.

It is hypothesised that the progression of cancer (encoded by the target variable *y*) can be modelled as a *causal kinetic model*. A *causal kinetic model* (originally proposed by [33]) extends the concept of *structural causal models* (*SCMs*), a popular approach to causal modelling, to the setting of *ordinary differential equations* (*ODEs*). While the *SCM* of a random variable *y* defines such a variable as a function of their causal parents (direct causes), the *causal kinetic model* assumes the form

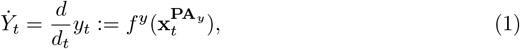

where 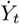 is the rate of change of the target variable, 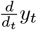 denotes the derivative of *y* (a target variable) at time *t*, and 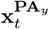 are the values of the set of parents of *y* at time *t*. In the proposed setting, it is assumed that the model of a target variable *y* (e.g., a biomarker gene), whose dynamics have changed during cancer due to the influence of driver genes, adopts the form shown in Eq. (1). Consequently, it is hypothesised that the set of parents of *y* (namely **PA**_*y*_) are cancer drivers.

#### 2.1.2 Invariance of structure of dynamic systems for causal discovery of cancer driver genes

One of the most notable properties of a causal system is that over the time that the causal structure holds, causal relationships remain *invariant* even under different interventions and/or environments [31, 33]. This *invariance* property is reflected, for example, in the fact that the conditional distribution of a variable *y* given its parents **PA**_*y*_ remains the same under interventions on variables other than *y*. One of the benefits of the *invariance* property is that it can be explored to assess causality.

In the proposed method, given a target variable *y* with dynamics that are influenced by cancer drivers, it is assumed that observed values of *y* have been generated by a causal model that is *invariant* during cancer progression. As the dynamics of *y* is influenced by cancer drivers, the derivative of *y* can be modelled as a function of its causal parents (i.e.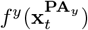), as shown in Eq. (1)). Therefore, the fitted values resulting from the evaluation of *f*^*y*^ will be reasonable estimations of the rate of change 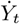 (from data) throughout the progression of the cancer when *f*^*y*^ is properly defined. This feature enables the dynamics of *y* to be used for identifying its causal parents.

Because gene expression data in cancer corresponds to real observations of the disease, it is possible to infer a function that describes the rate of change of a target variable *y* when a temporal dimension is included in the data. The *invariant* aspect of the model can be assessed by exploring cancer data collected from different “environments” (e.g. two repetitions of an experiment in which data on the same type of cancer are collected). As such data are uncommon in practice, the proposed method has the ability to emulate two different environments by systematically splitting the data (the details are in section 2.4). The data on the two environments are used to verify whether the model fits the values of the derivative estimated from the data on the target variable in both environments.

### 2.2 Method overview

The proposed method utilises cancer gene expression and the concept of *pseudotime* to emulate two environments and to create one pseudotemporally ordered data matrix for each environment. Subsequently, models of the form shown in Eq. (1) (the candidate causal structures) containing different combinations of putative drivers of the target *y* are tested in terms of their *invariance* across the environments. The better a model explains the data in both environments, the higher its ranking toward *invariance*. In other words, if a model provides consistent estimations of the derivative of *y* in both environments, then the proposed method ranks the model in a high position.

The outcome of the proposed method is a list of the top-ranked models as well as a list of cancer driver genes ranked in terms of their importance for those *invariant* models. The more times a gene appears in a highly ranked *invariant* model, the higher its rank as a cancer driver gene.

As shown in Fig. 1, the proposed method includes the following three phases:

**Figure 1.**
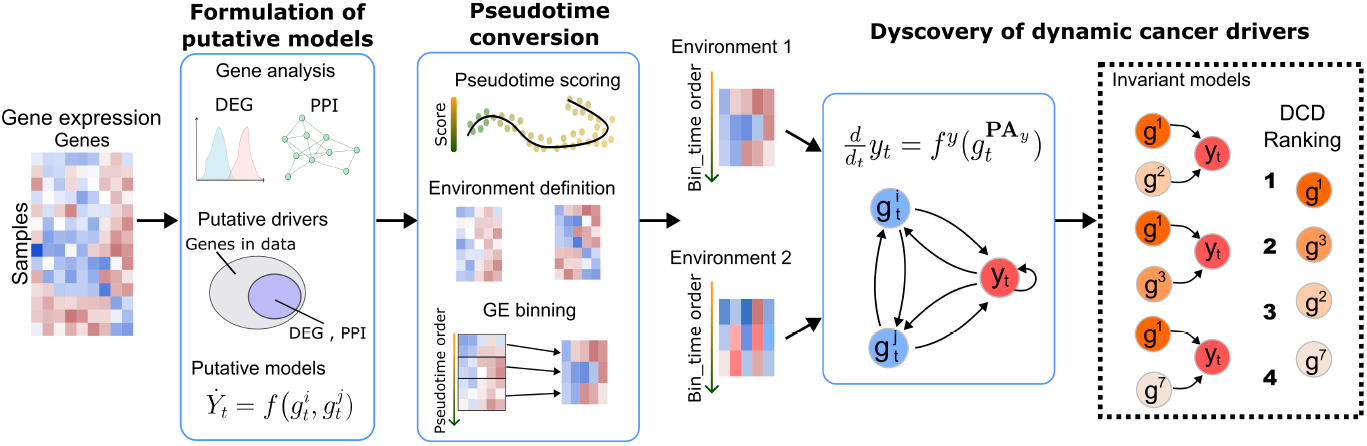
Summary of the proposed method for identifying cooperative genes that cause cancer progression. Starting with a gene expression dataset, the proposed method performs a putative model definition to create models that include only genes that have been selected as candidate cancer drivers by the method. Genes selected as putative cancer drivers are used for inferring a pseudotime order of samples that encode significant aspects of cancer progression. Ordered samples are grouped to form two mutually exclusive sub-datasets to emulate two different experimental environments. The gene expression and pseudotime score are binned to obtain Bin time series containing their expected values in each environment. Bin time series and the concept of invariance are used for finding the putative models that better describe the rate of change of a target variable *y* in both environments. Models that better explain the rate of change of *y* are considered invariant models. Genes in the invariant models are considered dynamic cancer drivers.

1. **Formulation of putative models**: A set of models (candidate causal structures) that describe the rate of change of the target variable *y* as a function of putative cancer drivers is created. Only genes with a reasonable potential of being cancer drivers (see section 2.3 for details) are kept in the putative models. These genes are selected by using differentially expressed gene (DEG) analysis and a protein-protein interaction (PPI) analysis.
2. **Pseudotime conversion**: Genes selected in the first phase are utilised for *pseudotime* scoring so that a temporal dimension is created and added into the data. Both the gene expression and *pseudotime* score are binned to emulate two environments and a single *pseudotime* (named *Bin time*).
3. **Discovery of dynamic cancer drivers**: A set of models are tested to observe how well they fit in both environments. The best fitted models are considered *stable towards invariance*, and genes consistently appearing in such models are selected as cancer driver genes (more details are in section 2.5). The results of this phase are a list of the dynamic models that are the most *stable towards invariance* as well as the set of cancer driver genes discovered in the data.

The details of the three phases are explained in the following sections.

### 2.3 Formulation of putative models

Gene expression is the result of a complex series of chemical reactions that can be modelled as random variables. Consequently, the *law of the mass action* (an essential principle in chemical reactions) is a reasonable selection of a model that fits the form in Eq. (1). This phase refers to the formulation of putative models (i.e. symbolic notation). Parameter estimations and model evaluations are performed in later phases.

In the proposed method, the parametric version of the *law of the mass action* [33] is adapted to the setup of the problem:

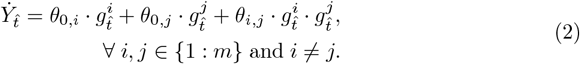

Specifically, models described by Eq. (2) correspond to the pairwise *main effect models* (as defined in [33]) of any pair of genes *g*^*i*^ and *g*^*j*^. In a *main effect model*, if the interaction term exists (i.e. *θ*_*i,j*_ ≠ 0), then the main effect terms also exist (i.e. *θ*_0,*i*_ ≠ 0, *θ*_0,*j*_ ≠ 0).

In addition to the pairwise *main effect models*, two lower order family models derived from Eq. (2) are included in the analyses: the pairwise gene collaboration linear models (i.e. *θ*_0,*i*_≠ 0, *θ*_0,*j*_≠ 0, and *θ*_*i,j*_ ≠ 0) and the pairwise interaction models (i.e. *θ*_0,*i*_ ≠ 0, *θ*_0,*j*_ ≠ 0, and *θ*_*i,j*_≠ 0).

In Eq. (2), 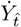 denotes the rate of change of the target variable, 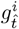 and 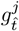 stand for the gene expression of genes *g*^*i*^ and *g*^*j*^ at 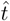, respectively, 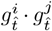 represents the interaction between *g*^*i*^ and *g*^*j*^ at 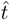, and *θ*_0,*i*_, *θ*_0,*j*_, and *θ*_*I,j*_ are model parameters. In this notation, 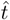 is explicitly used (instead of *t*) to highlight the fact that the parameter estimation does not use real time but a pseudotemporal reference (see section 2.4).

The analysis of gene interactions in data falls under the category of large scale system analysis. Ideally, a comparison among all possible combinations of different genes should be performed. However, a full exploration of the search space is infeasible due to the combinatorial nature of the problem.

To better illustrate this problem, a setup with only three genes of interest {*g*_1_, *g*_2_, *g*_3_} is assumed. Specifically in this setup, nine models are tested: the three pairwise *main effect models* (i.e. models with parameters [*θ*_0,1_, *θ*_0,2_, *θ*_1,2_],[*θ*_0,1_, *θ*_0,3_, *θ*_1,3_], and [*θ*_0,2_, *θ*_0,3_, *θ*_2,3_]), the three pairwise linear models (i.e. models with parameters [*θ*_0,1_, *θ*_0,2_], [*θ*_0,1_, *θ*_0,3_], and [*θ*_0,1_, *θ*_0,3_]), and the three pairwise interaction models (i.e. models with parameters [*θ*_1,2_], [*θ*_1,3_], [*θ*_2,3_]). Following a similar analysis, the search space expands to 18 models when considering four genes (as there are six pairwise *main effect models* in that scenario).

In general, there are 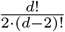 *main effect models* with the form shown in Eq. (2). As a result, the search space to be explored by the proposed method (that also includes the two lower-order family models) would consist of 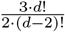 non-null models. This easily becomes infeasible for a large number of genes. Furthermore, models representing real driver gene interactions are sparse within the search space, as a sparsity of such interactions is expected. Accordingly, in this phase the number of models to be explored is reduced by including only genes that have a reasonable potential of being cancer driver genes.

In the proposed method, a gene has reasonable potential to be a cancer driver if it fulfills the following requirements:

1. The gene is differentially expressed between two meaningful conditions (such as normal vs cancer) in the dataset. This requirement is considered as fulfilled if such a gene is detected as differentially expressed after a DEG analysis.
2. The gene has the “potential” of influencing other genes. This requirement is considered as fulfilled if such a gene belongs to the PPI network. A cutoff of the degree of the gene (understood as the number of its interactions in the PPI network) can be used to further reduce the number of predictors when needed. The larger the degree, the more influential the gene.

Satisfying these requirements leads to a dimensionality reduction for the gene expression matrix from **G ℝ** ∈ ^*n*×*m*^ to **G** ∈ ℝ^*n*×*d*^, where *d* is the number of putative drivers. Using the *d* putative drivers as predictors, the family of models to be analysed by the proposed method is defined as shown in Eq. (2).

These considerations entail a significant reduction in the number of models to be explored while maintaining the predictive power of the proposed method.

### 2.4 Pseudotime conversion

This phase takes as input the dimensionally reduced gene expression matrix **G** ∈ **ℝ**^*n*×*d*^. This gene expression matrix is used for finding a *pseudotime* scoring that reflects the relative position of the samples throughout cancer progression. In the experiments, *Phenopath* [4] was used to calculate such a *pseudotime* score. *Phenopath* was selected because it is (to the best of our knowledge) the only *pseudotime* method explicitly designed for both single cell and bulk data.

Inferred *pseudotime* is used for the emulation of two environments (this is required in the proposed method for testing the *invariance* property). The two environments are emulated as follows:

1. Use **G** to find a *pseudotime* from the data.
2. Reorder the samples in **G** to follow that *pseudotime* in ascending order.
3. Split the ordered dataset by keeping the samples indexed by odd numbers in environment 1 and the samples indexed by even numbers in environment 2.
4. Split the *pseudotime* vector (ordered in ascending order) by keeping the values indexed by odd numbers as the *pseudotime* for environment 1 and the samples indexed by even numbers as the *pseudotime* for environment 2.

This heuristic, rather than random sampling, is used to split the dataset to maintain samples that reflect a similar disease progression in both environments.

At this stage, each environment has one half of the original samples and its corresponding *pseudotime* vector, with both following *pseudotime* ascending order. In the next step in this phase, the proposed method calculates the “binned version” of both the gene expression and *pseudotime* in each environment. To find the “binned version” of a vector (including the *pseudotime* vector and the gene vectors) in one environment, the following steps are performed:

1. Split the vector using the equal frequency method with a user-defined number of bins.
2. Map each bin into its expected value (i.e. mean).

This process is applied to all genes and *pseudotime* vectors in both environments. The result of this step consists of two matrices containing the “binned version” of genes, as well as the “binned version” of each environment’s *pseudotime*. Such matrices resemble the temporal progression of the disease because “binned versions” are created from samples ordered by following the *pseudotime*.

Finally, a common temporal reference for both environments is created by calculating the element-wise average of both *pseudotime* vectors. Hereafter, this common temporal reference is called *Bin time* (notated as 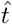 in Eq. (2)). As a result of this process, each environment has its own *Bin time* series for the gene expression of each gene and the target, with a common temporal reference (i.e. the *Bin time* vector).

### 2.5 Discovery of dynamic cancer drivers

The *Bin time* series obtained from the *pseudotime* conversion phase are used for assessing the stability of the models described in Eq. (2) in terms of their *invariance* across both environments. Testing whether a model is stable towards *invariance* can be achieved by testing if the estimations of the derivative of the target variable 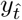 calculated from the observed values of *y* can be adequately fitted by models with the form shown in Eq. (2) [32].

Such stability towards *invariance* can be assessed by the *stability score* of the model [32]. The *stability score* is a goodness-of-fit metric that uses the residual sum of the squares of a fitted spline as an indicator of *non-invariance*. Moreover, this score can further guide a ranking procedure for the predictors (a detailed proof of the validity of both can be found in [33]). In the proposed method, the following steps are performed to calculate the *stability score* of a model *M* that has the form of the Eq. (2):

1. Fit a solution to the model *M* by using the data for the target and the predictors from environment 2. The *integrated model* described in [33] is replicated. This model implements the *trapezoidal rule* (a popular technique for approximating definite integrals) to obtain a numerical approximation of the model *M* as a function of the integral of its predictors. This approach is usually preferred over direct estimations of the derivative, as integration is numerically more stable than differentiation [7, 32]
2. Use the model parameters obtained from environment 2 to estimate the values of the derivative of the target in environment 1 by using the data for the predictors in the same environment (i.e. environment 1).
3. Fit a spline by using the data for the target in environment 1 as input and the estimated values of the target’s derivative (step 2) as constraints. The function *constrained*.*smoothspline* from the *CausalKinetiX* package [32] was used to fit such a spline. The residuals of the spline are expected to be small when the parameters found from environment 2 allow for an adequate fit of the spline for environment 1.

This procedure is performed a second time, but this time the order of the environments are interchanged. As a result, the proposed method creates two splines, one for environment 1 with the derivative constraint from environment 2, and one for environment 2 with the derivative constraint from environment 1. The mean of the residual sum of the squares of such splines is used as the *stability score*. The better the model fits in both environments, the lower its score. Consequently, in the proposed approach, models with low scores are considered to have a more stable fit towards *invariance*. Using this score leads to a ranking in which models with lower scores are better ranked. Such an inferred ranking is used for identifying cancer driver genes.

In practice, only a small fraction of the models in the search space are expected to be *invariant*. For this reason, it is suggested to consider as a starting point the top *l* = *d* models (where *d* is the number of putative drivers) as models stable towards *invariance*. Using this cutoff, a gene *g*^*i*^ can be scored to reflect the proportion of stable models that depend on such a gene, as described by

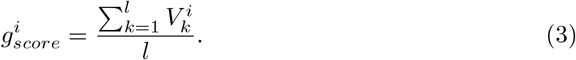

In particular, one can create the binary flag 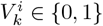 to specify whether or not the model *M*_*k*_ depends on the gene 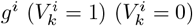. This score is used as an indicator of the causal impact that a gene has on cancer. Lowering the cutoff *l* makes the scoring procedure more strict (as the number of models considered decreases, the number of genes that appear in at least one invariant model also decreases). This potentially improves the performance of the method for certain datasets at the cost of an increase in the number of ties in a driver’s scores. Genes that consistently appear in the *invariant* models have higher scores and are considered cancer drivers by the proposed method.

The outcome of the proposed method includes a list of inferred cancer drivers ranked by their contribution toward the stability of the *invariant* models as well as the structure of the top *l* most stable invariant models. These invariant models contain (in general) interactions (interaction terms) and collaborations (main effect terms) of genes, as described by Eq. (2). Using this information, the proposed method creates a *gene collaborative network* by using any two genes in the invariant models as nodes and creating a link between genes that appear simultaneously in the same invariant model.

## 3 Experiments and results

### 3.1 Experimental methodology overview

In this section, we provide an intuitive overview of the key components of our methodology for identifying cooperative cancer drivers using dynamic causal inference.

- **Step 1: Data Preparation** We use gene expression data from both single cell and bulk sequencing datasets. Each dataset is preprocessed to normalise expression levels and remove low expressed genes. The pseudotime ordering is derived from this data, simulating a temporal trajectory of gene activity. This allows us to observe gene expression changes over an inferred timeline, rather than relying on static snapshots.
- **Step 2: Dynamic Causal Inference** We model the relationships between genes using causal kinetic models (CKMs). Unlike traditional correlation-based methods, CKMs account for temporal dependencies and causal interactions between genes. For example, if gene A consistently activates gene B in different experimental conditions (environments), we infer a causal relationship instead of just correlation.
- **Step 3: Cooperative Network Identification Using Causal Models** Our cancer driver identification is driven by causal inference, where we focus on identifying gene networks that have the highest impact on cancer progression. The *Stability score* quantifies the robustness of each causal model by assessing its *invariance property*, which is then used to rank potential drivers. When two genes appear in the same invariant model, they are connected within the network, highlighting their collaborative influence on the behaviour of other critical genes (our targets). This methodology prioritises genes that not only exhibit abnormal behaviour but also consistently work together to drive cancer progression.
- **Step 4: Validation** We validate the identified cooperative networks by cross-referencing our results with known cancer driver genes in existing literature. Additionally, we use statistical methods such as hyper-geometric testing to measure the enrichment of our identified genes among known cancer drivers, ensuring robustness and relevance.

### 3.2 Cancer drivers discovered from TCGA breast cancer data

Bulk data account for the vast majority of cancer data. Hence, experiments using the TCGA-BRCA dataset [46] were performed. The dataset was accessed in June 2022 and it was downloaded using the *TCGAbiolinks* package in R [13]. The dataset used in the experiments, containing gene expressions harmonised against GRCh38 (hg38), included 23694 genes from 1219 samples (113 normal tissue and 1106 primary tumour).

As part of the requirements of the proposed method, a target variable *y* with dynamics that have been altered during cancer needs to be provided. Well-known biomarkers or any gene with a gene expression that is altered because of cancer are suitable target variables provided that such alterations are captured in the analysed dataset.

In the experiments, the 40 genes identified by [30] (see Table 1) were explored as possible target variables. These genes were identified as influential in breast cancer based on data from 2433 primary breast cancer samples. A DEG analysis and a PPI analysis were performed to select the potential cancer drivers (i.e. the predictor genes in the models). For these experiments, a DEG analysis between normal and cancerous conditions was performed by using the function *TCGAanalyze_DEA* from the *TCGABiolinks* package (fdr.cut = 0.01, logFC.cut = 1, pipeline = “edgeR”, and method = “glmLRT”).

**Table 1.** **List of the 40 Mut-driver genes used in the experiments as target variables**, from [30].

|  |  |  |  |  |
| --- | --- | --- | --- | --- |
| SMAD4 | KDM6A | CDH1 | AKT1 | ZFP36L1 |
| USP9X | BRCA1 | CDKN1B | TBX3 | GATA3 |
| FOXP1 | TP53 | PIK3CA | CHEK2 | MAP3K1 |
| MEN1 | ARID1A | KMT2C | CBFB | GPS2 |
| MLLT4 | NF1 | AGTR2 | NCOR1 | CTCF |
| TBL1XR1 | SF3B1 | FOXO3 | KRAS | CTNNA1 |
| ERBB2 | MAP2K4 | PTEN | CDKN2A | BAP1 |
| PIK3R1 | RB1 | BRCA2 | RUNX1 | PBRM1 |

To perform the PPI analysis, the PPI network provided in [43] was used. To reduce the computational burden, a minimum of five interactions in the PPI network were selected as the threshold. This means that the putative cancer drivers are the genes identified as DEGs with more than four interactions in the PPI network. By following this procedure, 732 predictors (i.e. potential cancer drivers) were obtained. In real dynamic systems a variable can depend on itself (e.g. self-regulation); similarly, in the experiments the target gene *y* was included as an additional predictor variable.

Consequently, 733 variables (i.e. one target + 732 predictors) and a search space of 804834 models (as in Eq. (2)) per target gene were explored. Only 39 out of the 40 possible targets were further explored because the expression of the gene AGTR2 (ENSG00000180772) was missing from the dataset.

*Pseudotime* conversion was performed by using the *pseudotime* score inferred by *Phenopath* [4]. The 732 genes selected in the previous phase were used for *pseudotime* scoring. *Phenopath* requires a “path covariate” as additional input to guide the *pseudotime* scoring process so that it can be used in bulk data. The gene expression of ERBB2 (HER2) was selected to fulfill *Phenopath’s* requirement because this gene is both a well-known breast cancer driver and breast cancer biomarker frequently used for diagnosis and prognosis. The binning process involved placing 12 consecutive samples into one bin (approximately 1% of the dataset samples) after creating the two environments. As a result, *Bin_time* series with 50 *Bin_time* points were obtained for each environment.

#### 3.2.1 Systematic detection of cancer drivers by the proposed method

Assessing the performance of methods for discovering cancer drivers is a difficult task because no gold standard is available, and thus the “false positive rate” is not indicative of the performance of a method. However, it is a common practice to assess a method’s ability to systematically detect driver genes by using the Cancer Gene Census (CGC) [39] as a conservative approximation of the ground truth. The underlying logic of this common approach is that a method identifying more CGC genes from data than that expected by random selection (significance level ≤ 0.05) is less prone to misidentifications [42]. In the experiments, the hypergeometric test was used to evaluate whether CGC genes (CGC v94 n = 719 genes) can be systematically identified by using the driver score (Eq. (3)) inferred by the proposed method.

The hypergeometric test p.value was calculated by using the *phyper* function in R (p.value = 1 - *phyper(q - 1,m,n,k)*). The parameters of the test were as follows: *q* = number of CGC genes among the inferred cancer driver genes, *m* = number of CGC genes among the 733 model variables (baseline), *n* = number of non-CGC genes among the putative drivers, and *k* = number of inferred cancer drivers. The proposed method detects genes acting together (i.e. interacting and/or collaborating) to drive cancer progression rather than single gene drivers. Consequently, the hypergeometric test is used as an indicator of the systematic performance of the method as a whole, not as a performance metric of the method?s ability to detect single drivers nor of single experiments (i.e. a single target variable).

The inferences of the proposed method have a statistically significant overlap with the CGC gene set (p.value ≤ 0.05) when using the 11 target genes shown in Table 2. Furthermore, a significant overlap with the CGC set was found for seven additional targets when using the driver score (Eq. (3)) as the metric to select the top 150 inferred genes (see Table 3). These results, as well as the fact that the inferred sets of cancer drivers have relatively small sizes, suggest that the proposed method is able to effectively detect and rank cancer driver genes from bulk data. The results of the hypergeometric test for the 39 genes can be found online at https://github.com/AndresMCB/DynamicCancerDriverKM in *“supp_Table1 - hypertest for 39 target genes(Bulk)*.*csv”* (in the *supplementary* folder).

**Table 2.** Summary of the 11 target genes with a significant overlap with the CGC gene set. In the proposed method, the hypergeometric p.value is used as an indicator of systematic identification of cancer driver genes. The p.values were calculated by using the *phyper* function in R.

| Target $y$ | CGC baseline | n inferred | CGC detected | p.value |
| --- | --- | --- | --- | --- |
| MEN1 | 94 | 351 | 56 | 0.0101 |
| AFDN | 94 | 356 | 54 | 0.0413 |
| PIK3R1 | 93 | 445 | 69 | 0.0027 |
| TP53 | 94 | 277 | 44 | 0.0355 |
| NF1 | 94 | 317 | 50 | 0.0246 |
| PIK3CA | 94 | 351 | 53 | 0.0488 |
| FOXO3 | 94 | 367 | 56 | 0.0309 |
| BRCA2 | 93 | 327 | 53 | 0.0073 |
| CHEK2 | 94 | 278 | 45 | 0.0228 |
| CBFB | 94 | 282 | 44 | 0.0488 |
| CDKN2A | 93 | 198 | 38 | 0.0013 |

**Table 3.** Summary of the seven additional targets with a statistically significant overlap with the CGC gene list when using the top 150 genes ranked by the proposed method. The p.value resulting from the hypergeometric test is shown in the last column. The p.values were calculated by using the *phyper* function in R.

| Target *y* | CGC baseline | CGC in top 150 | p.value |
| --- | --- | --- | --- |
| SMAD4 | 94 | 27 | 0.0261 |
| FOXP1 | 94 | 26 | 0.0462 |
| CDH1 | 94 | 30 | 0.0034 |
| KMT2C | 94 | 30 | 0.0034 |
| NCOR1 | 94 | 27 | 0.0261 |
| CTNNA1 | 93 | 29 | 0.005 |
| PBRM1 | 94 | 27 | 0.0261 |

Besides the ability to systematically detect driver genes, an effective cancer driver discovery method is expected to retrieve genes that are strongly linked to cancer in biological terms. To assess whether the inferred cancer drivers are enriched in terms of cancer, the following analyses were performed via the *Enrichr* online tool [48]: *KEGG 2021 Human* (biological pathways), *GO Biological Processes 2021*, and *DisGeNET* (human diseases). The experiments for the targets PIK3R1 and CDKN2A were selected for these analyses, as they have the largest (445 genes) and the smallest (198 genes) inferred sets, respectively, and significantly overlap with the CGC list.

In both cases, the inferred set was significantly enriched with respect to cancer. For both analysed targets, the top one *KEGG* enriched term (ranked by p.value) was **pathways in cancer** (PIK3R1set p.val = 3.609e-27, CDKN2Aset p.val = 1.544e-15), and all other terms in the top four were pathways related to cancer, including **focal adhesion** (PIK3R1set p.val = 1.464e-16, CDKN2Aset p.val = 1.609e-12), **transcriptional misregulation in cancer** (PIK3R1set p.val = 3.428e-16), and **proteoglycans in cancer** (CDKN2Aset p.val = 2.297e-10).

Similarly, all top five *DisGeNET* and all top five *GO Biological Processes* enriched terms (ranked by p.value) for both inferred sets were strongly related to cancer. The top five *DisGeNET* enriched terms included **neoplasm metastasis** (PIK3R1set p.val = 3.366e-63, CDKN2Aset p.val = 2.894e-50), **breast carcinoma** (PIK3R1set p.val = 4.476e-64, CDKN2Aset p.val = 8.765e-47), and **mammary neoplasms** (PIK3R1set p.val = 3.366e-63, CDKN2Aset p.val = 3.720e-54). The top five *GO Biological Processes* enriched terms include **positive regulation of transcription**,

**DNA-templated (GO: 0045893)** (PIK3R1set p.val = 5.884e-28, CDKN2Aset p.val = 1.213e-16), and **positive regulation of transcription by RNA polymerase II (GO:0045944)** (PIK3R1set p.val = 1.479e-24, CDKN2Aset p.val = 1.144e-15). The complete results for the enrichment analyses using bulk data can be found online at https://github.com/AndresMCB/DynamicCancerDriverKM in the *supplementary* folder (*supplementary tables 3-8*).

### 3.3 Identification of mutated and non-mutated cancer drivers by the proposed method

For this experiment, the single nucleotide variant (SNV) frequency was included in the set of inferred cancer drivers to assess the ability of the proposed method to identify cancer drivers regardless of their mutational status. This is one of the key aspects of the proposed method, as not all cancer driver genes are mutated.

Because non-mutated cancer drivers have been investigated minimally in the literature, it is difficult to evaluate the performance of non-mutated driver inference methods. Thus, this analysis was restricted to highly reliable driver genes inferred by the proposed method, as those consistently detected in many experiments were less likely to be false positives. Specifically, only the genes that were detected as cancer drivers in more than 50% of the 11 experiments summarised in Table 2 were considered.

The SNV frequency was incorporated into the resulting set, which contained 308 genes. Mutational information was obtained from the TCGA-BRCA dataset in Mutation Annotation Format (MAF) (MAF.hg38) by using the R package *TCGAbiolinks*.

The SNV frequency was used to classify the cancer driver discoveries as mutated (n=280) or non-mutated (n=28). The Network of Cancer Genes & Healthy Drivers (NCG 7.0) was used to further analyse the top 10 genes with the highest SNV frequency and all non-mutated genes [18]. The NCG is a manually curated collection of cancer genes that contains canonical cancer drivers (such as the CGC genes) and highly confident candidate drivers.

As expected, the proposed method was able to consistently recover canonical/candidate cancer drivers, including non-mutated drivers (see Table 4). Seven out of the top 10 most mutated inferred genes (namely SYNE1, DST, DMD, CUBN, ANK2, LRP1, and F8) are NCG genes while PTPRB is a CGC gene with the role of a tumour suppressor. The genes CALR, ALDH2, and CDKN2A are CGC genes identified by the proposed method that have no mutation in the TCGA-BRCA dataset.

**Table 4.**
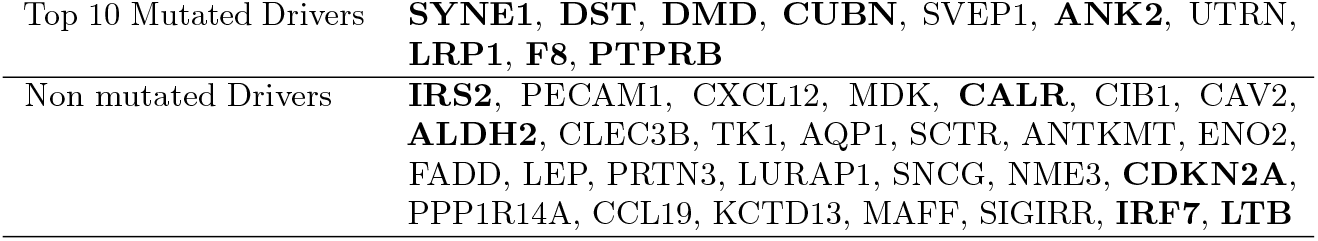
Top 10 most mutated (SNV) and all non-mutated inferred cancer drivers in the intersection set of the results for the 11 target genes shown in Table 2. Only genes identified in more than 50% of the 11 experiments (i.e. 6 or more) were considered. Canonical (CGC) and highly confident drivers (NCG) are shown in boldface font. The NCG7.0 analysis was performed using the online tool at http://ncg.kcl.ac.uk/.

|  |  |
| --- | --- |
| Top 10 Mutated Drivers | **SYNE1, DST, DMD, CUBN, SVEP1, ANK2, UTRN, LRP1, F8, PTPRB** |
| Non mutated Drivers | **IRS2, PECAM1, CXCL12, MDK, CALR, CIB1, CAV2, ALDH2, CLEC3B, TK1, AQP1, SCTR, ANTKMT, ENO2, FADD, LEP, PRTN3, LURAP1, SNCG, NME3, CDKN2A, PPP1R14A, CCL19, KCTD13, MAFF, SIGIRR, IRF7, LTB** |

Additionally, the proposed method identified three other highly confident drivers (i.e. IRS2, IRF7, LBT) that have no mutation in the analysed dataset. To name a few of the novel non-mutated cancer drivers discovered by the proposed method, CXCL12 (specifically the CXCL12/CXCR4 signal pathway) has been found to promote several aspects of breast cancer tumorigenesis, such as proliferation, cell motility, and distant metastasis [22, 36, 52]. TK1 and MDK have also been linked to advanced/metastatic stages of breast cancer [16, 17, 19, 23, 51], while CIB1 is a promising therapeutic target gene that drives cancer cell death [8, 9]. The relevance of the discovered genes to cancer provides strong support for the effectiveness of the proposed method in identifying mutated and non-mutated drivers of cancer.

### 3.4 Inferred cancer driver sets from single cell data

To assess the functionality of the proposed method for single cell data, the single cell RNA sequencing data from the NCBI GEO database (accession GSE75688) [10] was analysed. The original dataset contains gene expressions of 515 single cells from 11 breast cancer patients with distinct molecular subtypes: oestrogen receptor positive (ER+), human epidermal growth factor receptor 2 positive (HER2+), double positive (ER+ and HER2+), and triple-negative breast cancer (TNBC). Out of 515 cells in this dataset, 317 correspond to tumour cells while the other 198 correspond to immune (n=175) and stromal (n=23) cells. After removing low expressed genes, the dataset had 9288 genes.

For these experiments, the putative model formulation phase was performed by including differentially expressed genes with a node degree of one in the PPI network (i.e. genes with at least one interaction in the PPI). A DEG analysis (fdr.cut = 0.01, logFC.cut = 1, pipeline = “edgeR”, and method = “glmLRT”) between the cancerous and normal conditions was performed by using the single cells labelled as “tumour” for cancer and the cells labelled as “stromal” for normal. This procedure led to the selection of 178 genes for further exploration. A total of 340 samples were used for the data-driven inference process (tumour and stromal cells). For the *pseudotime* conversion phase, the number of *Bin_time* points was set to 30 to allow at least five samples per *Bin_time* point during the binning process.

The 39 Mut-driver genes in Table 1 that are present in the dataset were used as possible targets (AFDN was missing). The results for all 39 genes were similar in terms of the size of the inferred sets (84-129, mean = 105, sd = 9.11) and the size of the intersection with the CGC set (12-19, mean = 14.79, sd = 1.67). The proposed method’s inferences showed a significant overlap with the CGC gene set for seven of the explored target genes (SMAD4, RB1, CDKN1B, FOX03, PTEN, ZFP36l1, and MAP3K1) when considering all genes with a score (as calculated by using Eq. (3)) greater than zero. The number of targets with a significant overlap increased to 14 (see Table 5) when considering the top 80 genes of each target (selected as the cutoff because the size of the smaller inferred set among the single cell experiments was 84).

**Table 5.**
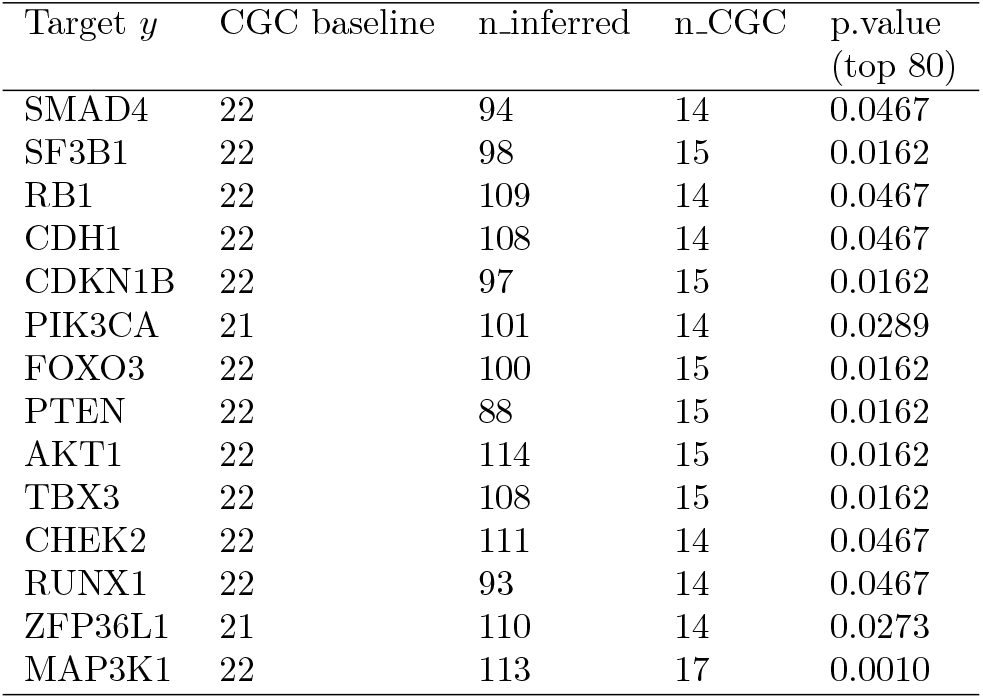
Summary of single cell data experiments for the 14 target genes with a significant overlap with the CGC gene set when considering the top 80 inferred genes per target. The p.values were calculated by using the *phyper* function in R.

| Target *y* | CGC baseline | n_inferred | n_CGC | p.value<br>(top 80) |
| --- | --- | --- | --- | --- |
| SMAD4 | 22 | 94 | 14 | 0.0467 |
| SF3B1 | 22 | 98 | 15 | 0.0162 |
| RB1 | 22 | 109 | 14 | 0.0467 |
| CDH1 | 22 | 108 | 14 | 0.0467 |
| CDKN1B | 22 | 97 | 15 | 0.0162 |
| PIK3CA | 21 | 101 | 14 | 0.0289 |
| FOXO3 | 22 | 100 | 15 | 0.0162 |
| PTEN | 22 | 88 | 15 | 0.0162 |
| AKT1 | 22 | 114 | 15 | 0.0162 |
| TBX3 | 22 | 108 | 15 | 0.0162 |
| CHEK2 | 22 | 111 | 14 | 0.0467 |
| RUNX1 | 22 | 93 | 14 | 0.0467 |
| ZFP36L1 | 21 | 110 | 14 | 0.0273 |
| MAP3K1 | 22 | 113 | 17 | 0.0010 |

The inferred driver sets from those 14 targets were further explored to assemble a single driver list. Table 6 shows the cancer driver genes inferred from the single cell data that were detected by the proposed method for at least 10 different target genes (i.e. they appear in the intersection of at least 10 of the inferred sets). The genes ATP1B1, EZR, FBLN1, HLA-E, and TAPBP were identified as cancer drivers by the proposed method in all 14 targets, suggesting these genes are highly relevant to cancer progression for the 11 patients of the study. Among those five genes, EZR is a CGC gene, while ATP1B1 and FBLN1 are designated as *mouse genes* in the *Catalogue Of Somatic Mutations In Cancer (COSMIC)* [41]. A *COSMIC mouse gene* is a gene with experimental support (mouse insertional mutagenesis) that strongly suggests such a gene as a cancer-causing gene. The full list of inferred cancer drivers from the single cell data for each of the 39 experiments can be found at https://github.com/AndresMCB/DynamicCancerDriverKM in *“supp_Table9 - inferred drivers from single cell*.*xlsx”* (the *supplementary* folder).

**Table 6.** Cancer driver genes inferred from the single cell data that were detected by the proposed method in at least 10 different experiments (different target genes).

| n targets $y$ | Inferred drivers |
| --- | --- |
| 14 | ATP1B1, EZR, FBLN1, HLA-E, TAPBP |
| 13 | CALU, CD99, DHX9, DPYSL2, ITM2B, PDIA4, RNF19A, SERPING1, SNRPF, STOM, SYNE1, TPD52 |
| 12 | ARF1, AXL, CDK12, COL1A1, FLNA, GLRX3, GSN, HLA-C, IFI16, LRPPRC, PIK3CA, SND1, UBR5, VIM, ZNF24 |
| 11 | ABCA1, ACTR1B, BCCIP, C3, DAB2, ERBB2, FER, FOXP1, FTL, HSP90AB1, IGFBP4, IGFBP7, ITGB1, LAMP1, MUC1, OCLN, PRSS23, PTPRF, RIN2, RPS6KA3, SIX1, STAT1, STAT2, TNPO2, TOB1, WWTR1 |
| 10 | ANXA5, B2M, DNAJC10, ERBB3, GLI3, IKZF5, JPT2, PEX13, PTGFRN, TAX1BP1, TCF4, TFAP2A, TFAP2C, TGFB3, UBA1 |

As the proposed method detects genes that act together (i.e., interact and/or collaborate) to drive cancer progression rather than identifying single gene drivers, the group gene cooperation networks obtained from the experiments were examined.

Specifically, the networks inferred when using PTEN and AKT1 as target variables (hereafter referred to as **PTENnetSC** and **AKT1netSC**, respectively) were explored. These two experiments were chosen because they returned the smallest and largest sets of inferred cancer drivers, respectively. A community detection process was performed by using the function *group_infomap* from the *tidygraph* R package [29] and using the default parameters (see Fig. 2).

**Figure 2.**
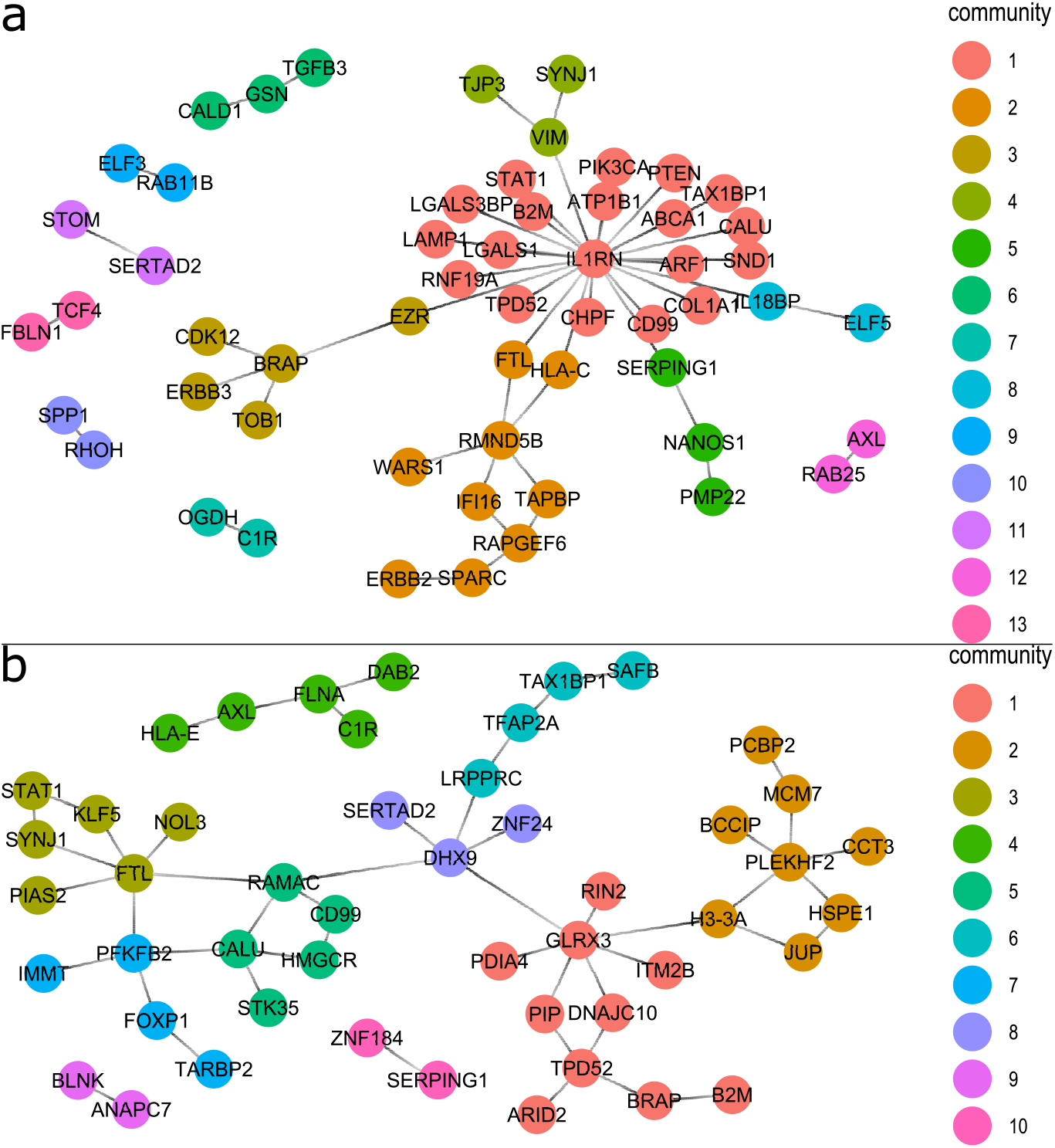
Inferred gene cooperation networks when using the top 50 models of the experiments with a) PTEN (PTENnetSC) and b) AKT1 (AKT1netSC) as target variables.

For the sake of visualisation, the networks were built by only considering the top 50 models per target ranked by their *stability score* (as described in section 2.5). The network nodes correspond to the genes appearing in any of the top *invariant* models.

The edges in the networks link genes appearing simultaneously in at least one of the top *invariant* models. As expected, the inferred networks included canonical driver genes. Specifically, FLNA, B2M, H3-3A, ARID2, and FOXP1 in **AKT1netSC**, and COL1A1, ERBB3, CDK12, PIK3CA, EZR, SND1, ELF3, PTEN, B2M, ERBB2, and RHOH in **PTENnetSC**.

### 3.5 Biological Interpretation of Identified Gene Networks

Our analysis from Bulk data identified several novel gene candidates, such as CXCL12 and TK1, that appear to play key roles in cancer progression. CXCL12 is a chemokine known for its role in the tumour microenvironment, where it influences cancer cell migration and invasion. Its identification as a potential driver gene in our model suggests that it may have broader implications in regulating the network of genes associated with cancer progression.

Likewise, TK1, a thymidine kinase involved in DNA synthesis, has been implicated in cell proliferation and tumour growth. Our analysis positions TK1 as a hub gene, with multiple downstream targets influenced by its activity. This insight is significant, as it suggests that TK1 may be a central regulator of gene networks driving unchecked cell division in cancer.

To further validate these findings, we examined the pathways in which these genes are involved. Both CXCL12 and TK1 were found to be enriched in key oncogenic pathways, such as the PI3K/AKT pathway, which is known to promote cell survival and growth. These findings align with previous research, reinforcing the biological plausibility of our results.

Similarly, from our analyses on single cell data, the inferred novel drivers FTL, DHX9, GLRX3, and PLEKHF2 were central nodes in **AKT1netSC**, while IL1RN and RMND5B were central nodes in **PTENnetSC**. FTL, PLEKHF2, and DHX9 are

*COSMIC mouse genes*, and DHX9 is also a putative oncogene in the NCG 7.0 database. Additionally, it has been confirmed that IL1RN is a tumour-associated gene that is negatively correlated with the survival time of glioma patients [44], and it has also been found to be upregulated across the ephithelial-to-mesenchymal transition (EMT) in triple negative breast cancer [1].

Finally, the community surrounding the IL1RN gene (i.e. community 1 in Fig. 2-a) was explored, as it was the largest community detected among these two networks (19 genes). A series of pathway analyses was performed to verify the biological relevance of the inferred cluster. *Reactome 2022, KEGG 2021 Human*, and *Elservier Pathway collection* enrichment analyses were performed via the *Enrichr* online tool [48]. Within the top five enriched terms of each analysis (ranked by combined score), four *Reactome 2022*, one *KEGG 2021 Human*, and four *Elservier Pathway collection* enriched terms were strongly linked to cancer (see Table 7).

**Table 7.** Enriched biological pathways for the IL1RN gene cluster (community 1 in Fig. 2a) in Reactome 2022, KEGG human 2021, and Elsevier Pathway Collection. The IL1RN inferred gene community was enriched in several biological pathways closely related to cancer. The enrichment analysis was performed by using Enrichr https://maayanlab.cloud/Enrichr/.

| Database | Term index | Enriched Terms | P.value | Combined Score |
| --- | --- | --- | --- | --- |
| Reactome 2022 | 1 | Signaling by PDGFRA Extracellular Domain Mutants R-HSA-9673770 | 0.000046 | 2602.87 |
|  | 2 | Signaling by Cytosolic FGFR1 Fusion Mutants R-HSA-1839117 | 0.000115 | 1420.01 |
|  | 3 | Signaling by KIT In Disease R-HSA-9669938 | 0.000144 | 1221.34 |
|  | 4 | Signaling by PDGFR In Disease R-HSA-9671555 | 0.000144 | 1221.34 |
| KEGG Human 2021 | 2 | PD-L1 expression and PD-1 checkpoint pathway in cancer | 0.000078 | 410.07 |
| Elsevier Pathway Collection | 1 | Genes with Mutations Associated with Hereditary Breast and/or Ovarian Cancer Syndrome | 0.000030 | 3488.77 |
|  | 2 | Genes with Mutations Associated with Endometrial Cancer | 0.000038 | 2987.15 |
|  | 3 | PI3K/AKT/MTOR Signaling Activation in Cancer | 0.000115 | 1420.01 |
|  | 5 | Genes with Mutations in Cancer Metabolic Reprogramming | 0.000144 | 1221.34 |

## 4 Conclusions

This paper presents a new causal inference method for cancer driver discovery, considering the dynamics of gene interactions and collaborations to identify sets of cancer drivers. The proposed method utilises the fact that driver genes alter the dynamics of core processes in cancer cells by dysregulating biological pathways. One interesting advantage of this method is that it relaxes the constraints of detecting mutational events by focusing on the observed changes in cancer driver expression and their causal influence on other genes during cancer development. This allows the proposed method to be used for identifying cancer drivers that do not have mutations.

Causal kinetic models of cancer drivers are considered an extension of structural causal models to the setting of ordinary differential equations. Assessing the causal effect of a driver gene on other genes, to distinguish a cancer driver (the cause) from affected genes (the effects), is a sensible approach to reduce spurious discoveries. The experiments suggest that the inclusion of formal procedures for testing the causal structure of the underlying dynamics of cancer helps infer driver genes strongly related to altered biological processes in cancer.

Another interesting aspect of the proposed method is that it produces models of gene interaction dynamics, which provide insight into the temporal behaviour of genes during cancer. Such models can be utilised to represent the interactions throughout cancer as a cooperative network. The network itself represents the set of drivers acting together (not necessarily regulatory relationships) to induce cancer progression. The inferred networks can be potentially used for identifying clusters of genes that dysregulate specific processes during cancer. As a result, the network provides insights into the mechanisms that may cause carcinogenesis and early cancer development.

While our study focuses on breast cancer, the principles underlying our method are not cancer-specific. The integration of dynamic causal inference and pseudotime analysis could be adapted to study other types of cancer or diseases where gene expression data is available. Future work could explore the application of this method to cancers with distinct genetic landscapes, such as lung or pancreatic cancer.

Additionally, adapting the method to diseases characterised by other forms of gene dysregulation, such as neurodegenerative diseases, could provide valuable insights into their progression. This flexibility highlights the potential for our approach to make a significant impact across multiple areas of biomedical research.

## 5 Acknowledgments

We express our gratitude to the anonymous reviewers for their insightful comments, which have significantly improved the quality of this manuscript. We also thank our colleagues and collaborators from the UniSA data analytics group who contributed to the development and validation of this method.

This work is supported in part by funds from the Australian Technology Network ATN-LATAM Research Scholarship Scheme, and Australian Research Council ARC - Discovery Early Career Researcher Award (DECRA, DE 200100200).

## 6 Author contributions statement

**Andres M. Cifuentes-Bernal**: Conceptualisation, Methodology, Software, Validation, Formal analysis, Investigation, Writing – Original Draft, Visualisation. **Thuc Duy Le**: Conceptualisation, Methodology, Validation, Formal analysis, Investigation, Writing – Original Draft, Supervision. **Lin Liu**: Writing - Review & Editing. **Jiuyong Li**: Writing - Review & Editing.

